# Unraveling the developmental heterogeneity within the human retina to reconstruct the continuity of retinal ganglion cell maturation and stage-specific intrinsic and extrinsic factors

**DOI:** 10.1101/2024.10.16.618776

**Authors:** Emil Kriukov, Jonathan R. Soucy, Everett Labrecque, Petr Baranov

## Abstract

Tissue development is a complex spatiotemporal process with multiple interdependent components. Anatomical, histological, sequencing, and evolutional strategies can be used to profile and explain tissue development from different perspectives. The introduction of scRNAseq methods and the computational tools allows to deconvolute developmental heterogeneity and draw a decomposed uniform map. In this manuscript, we decomposed the development of a human retina with a focus on the retinal ganglion cells (RGC). To increase the temporal resolution of retinal cell classes maturation state we assumed the working hypothesis that that maturation of retinal ganglion cells is a continuous, non-discrete process. We have assembled the scRNAseq atlas of human fetal retina from fetal week 8 to week 27 and applied the computational methods to unravel maturation heterogeneity into a uniform maturation track. We align RGC transcriptomes in pseudotime to map RGC developmental fate trajectories against the broader timeline of retinal development. Through this analysis, we identified the continuous maturation track of RGC and described the cell-intrinsic (DEGs, maturation gene profiles, regulons, transcriptional motifs) and -extrinsic profiles (neurotrophic receptors across maturation, cell-cell interactions) of different RGC maturation states. We described the genes involved in the retina and RGC maturation, including de novo RGC maturation drivers. We demonstrate the application of the human fetal retina atlas as a reference tool, allowing automated annotation and universal embedding of scRNAseq data. Altogether, our findings deepen the current knowledge of the retina and RGC maturation by bringing in the maturation dimension for the cell class vs. state analysis. We show how the pseudotime application contributes to developmental-oriented analyses, allowing to order the cells by their maturation state. This approach not only improves the downstream computational analysis but also provides a true maturation track transcriptomics profile.

## Introduction

The complexity of the mammalian retina arises from the unique combination of extrinsic and intrinsic factors in its development. Extensive lineage tracing, transcriptomic and transplantation studies confirm the paradigm of intrinsically different retinal progenitors (*1*), capable of producing different types of retinal neurons. The current paradigm, supported by anatomical, genomic, transcriptomic and evolutionary data, implies that the same holds true across all mammalian species (*2*), including humans (*3*). Several single-cell and single-nuclei RNA seq datasets have been generated(*4–10*). They allowed us to look into human retinal development with higher temporal resolution and approach the questions of cell fate specification, cell class and cell type heterogeneity at any given point in development, and regulation of cell trajectory by intrinsic and extrinsic factors.

Here, we establish a human fetal retinal atlas by integrating currently available human fetal retina single-cell RNA sequencing datasets(*5, 6*) into a single resource to create a high-resolution map of retinal development. We used advanced integrated transcriptomics methods, including pseudotime(*11*) with the cell fate trajectory reconstruction(*12*), potency(*13*), and trajectory density(*14*) methods to deconvolute the transcriptional signal.

This atlas highlighted the heterogeneity of retinal ganglion cells at every timepoint of ontogenesis and the ability of computational methods to reconstruct the development path and align individual cell transcriptomes to it. By doing so, we were able to visualize the continuity of RGC development with the exact transcriptional changes along the way. We also explored the regulome and interactome of human RGC throughout development, which allows us to separate intrinsic and extrinsic signals, respectively.

## Methods

### Single-cell RNA Sequencing Data Processing

Single-cell RNA sequencing data analysis was performed in RStudio v. 2023.06.1 Build 524 (R language v. 4.3.1-4.4.1) using previously published datasets from the human fetal retina (from Week 8 to Week 27)(*5, 6*). The raw Cell ranger output data was processed independently following the standard Seurat pipeline with a few differences: for FindNeighbors() and RunUMAP(), dims = 50. For multiple stages, the analysis included both the peripheral and central retina. Individual Seurat objects are stored separately in the data availability folder.

### Data Integration and Atlas Generation

To integrate the datasets upon independent processing of every object, we follow the integration pipeline from Seurat(*15*) v.4, which includes the steps of SelectIntegrationFeatures(), FindIntegrationAnchors(), IntegrateData(), and a custom script for the dataset processing in the same manner as for the independent datasets from the previous step.

To annotate the cell types, we manually annotated the data by identifying variable features for every Seurat cluster. This was later confirmed by integrating our data with the human brain cortex atlas from the Azimuth HubMAP source(*16*) to prove the neuronal identity using the prediction score. The annotated cell classes gave us insights into the type frequency information obtained by merging the time points within every 4-week step. The frequency for major retinal cell classes was calculated as several cells per cell type/class to the total number of cells in the dataset of the relevant time point. For further data analysis, the Azimuth approach and the AzimuthReference() and SCTransform()(*17*) functions can be used to use this atlas as a reference tool and build a representative UMAP embedding with automated cell type annotation. The detailed notebook is provided in the paper GitHub repository.

### Cell-cell interactions analysis

To obtain the information related to incoming and outgoing cluster ligand-receptors interactions, we used the CellChat package (v. 2.1.0) for every time point using our atlas as previously described (*18, 19*). To avoid the projection reducing the dropout effects of signaling genes, particularly for possible zero expression of subunits of ligands/receptors, we used raw.use = FALSE setting in computeCommunProb() function. Total heatmaps for the incoming and outgoing signaling and chord diagrams for the signaling of interest were generated. Additionally, we used LRLoop(*20*), following the default pipeline, to calculate the communication levels of every ligand-receptor pair in the package database per time point to build the patterns of communication across development. We labeled the retinal ganglion cells cluster as a cluster of interest while the rest of the populations were merged into one group.

### Python Data Transformation and ForceAtlas2-scFates pseudotime application

Once the atlas was generated, we converted it from.h5Seurat to .h5ad format for the Python-dependent downstream analysis (Python v. 3.7). We used scanpy(*21*) to generate ForceAtlas2-based(*22*) embedding for the RGC subset of the atlas with n_eigs = 3 and additional palantir-based(*23*) preprocessing. With the embedding to be generated, we reconstructed the cell fate trajectory using the scFates package(*11*). On that trajectory, pseudotime values were quantified, and RGC were split into t_factor groups with the pseudotime values order. With the in-built scFates functions, we found variable genes on the cell fate trajectory of RGC development. The metadata, including pseudotime and t_factor clustering, was exported to be added to the Seurat object with the AddMetaData() function in Seurat.

### Cell Fate Trajectory: RNA Velocity, CytoTRACE2, Mellon

We have applied a combination of methods supporting each other in understanding the cell fate trajectory. The analysis for this part was performed in R and Python.

For the R part, the master .h5Seurat object with the atlas was used to generate the potency results using CytoTRACE2(*13*). We followed the default pipeline for the full model described in the package.

The Python part included the downstream analysis of Cell ranger output. We applied velocyto(*12*) to generate cellsorted_possorted_bam.bam file from possorted_bam.bam file using the same reference genomes as the ones used in the public data integrated in the atlas. From the cellsorted_possorted_bam.bam file we generate .loom file and integrate it with the master .h5ad object using scvelo. Further downstream RNA Velocity(*12*) analysis was performed in scvelo(*24*), and the trajectory was built using dynamical mode.

The master.h5ad object was used to generate the cell density results by utilizing the mellon package following the default Mellon(*14*) pipeline.

### Downstream Analysis

The atlas with additional metadata was used for the further GSEA pathway analysis(*25*) in R using escape package(*26*) (v. 1.6.0) and ssGSEA method(*27*). We selected multiple pathways from the Molecular Signatures Database related to retinal ganglion cell function and performed the enrichment to visualize it with the canonical time points and t_factor (pseudotime groups) as metadata. For the GOBP_NEURAL_RETINA_DEVELOPMENT, GOBP_APOPTOTIC_PROCESS_INVOLVED_IN_DEVELOPMENT, GOBP_AXON_DEVELOPMENT, GOBP_AXON_EXTENSION, GOBP_NEURON_ MATURATION, and GOBP_NEURON_MIGRATION pathways, we obtained the gene list to build dot plots and ridge enrichment plots split by both canonical time points and t_factor clusters.

Furthermore, we established a list of neurotrophic ligands and their potential receptors from currently available ligand-receptor databases. We generated heatmaps and expression dynamics plots to show the trend of neurotrophic receptor expression across development (for both canonical time points and pseudotime).

### Data Availability Statement

The software and packages used for the analysis can be found in Table S1. The code used for this study to reproduce the bioinformatical analysis is available on GitHub (link available upon publication). The GitHub repository provides detailed instructions on every step of the analysis and the code to reproduce the computational analysis. To obtain the data generated in this study, please refer to the relevant repository folder or contact the corresponding authors. This data may include individual Seurat objects, integrated atlas, reference atlas version, Python cell fate reconstructed RGC subset, individual and integrated CellChat objects, and the gene/ligand-receptor lists generated throughout the analysis.

To make our discoveries more transparent and available, we deposited the atlas for the public audience on the CellxGene platform. This allows users to perform basic data analysis in the web browser. The atlas (current v.1) is available upon publication. Please contact the corresponding author for the Seurat and CellChat files in ‘.rds’ format.

## Results

### Establishing a Human Fetal Retinal Atlas to Study Extrinsic and Intrinsic Patterns of Retinal Cell Classes

We sourced, reanalyzed, and integrated published human fetal and adult retina single-cell RNA sequencing data to establish a human fetal retinal atlas(*5, 6*). The resulting atlas contains 108,838 cells from fetal weeks 8 to 27 with a one-week sample resolution. We keep the original fetal day resolution timepoint where possible. We performed cluster identification using single-cell RNA sequencing data to define each retinal population/subpopulation and major retinal classes: retinal ganglion (RGC), Bipolar, amacrine (AC), horizontal (HC), bipolar, rod, cone, Müller glia, and progenitor cells throughout development based on their cell type-specific genes (*SOX2, MKI67, PAX6, ATOH7, VIM, RLBP1, OTX2, PDE6H, RHO, etc.)* (**Figure 1A-B**). As a resource to the community, we also described the complete dynamics of regulons (**Figure S1-2**), transcriptional motifs, and cell-cell interactions within and in-between identified cell populations (**Figure S3**) in the resolution available using LRLoop, CellChat (*18, 19*), and pySCENIC (*28*). Within the resolution, we cover every cell class information on the currently available ligand-receptor database in CellChat v. 2 (*19*) and regulons and motifs from the cisTarget 2017 motif collection database for hg19.

**Figure 1.**
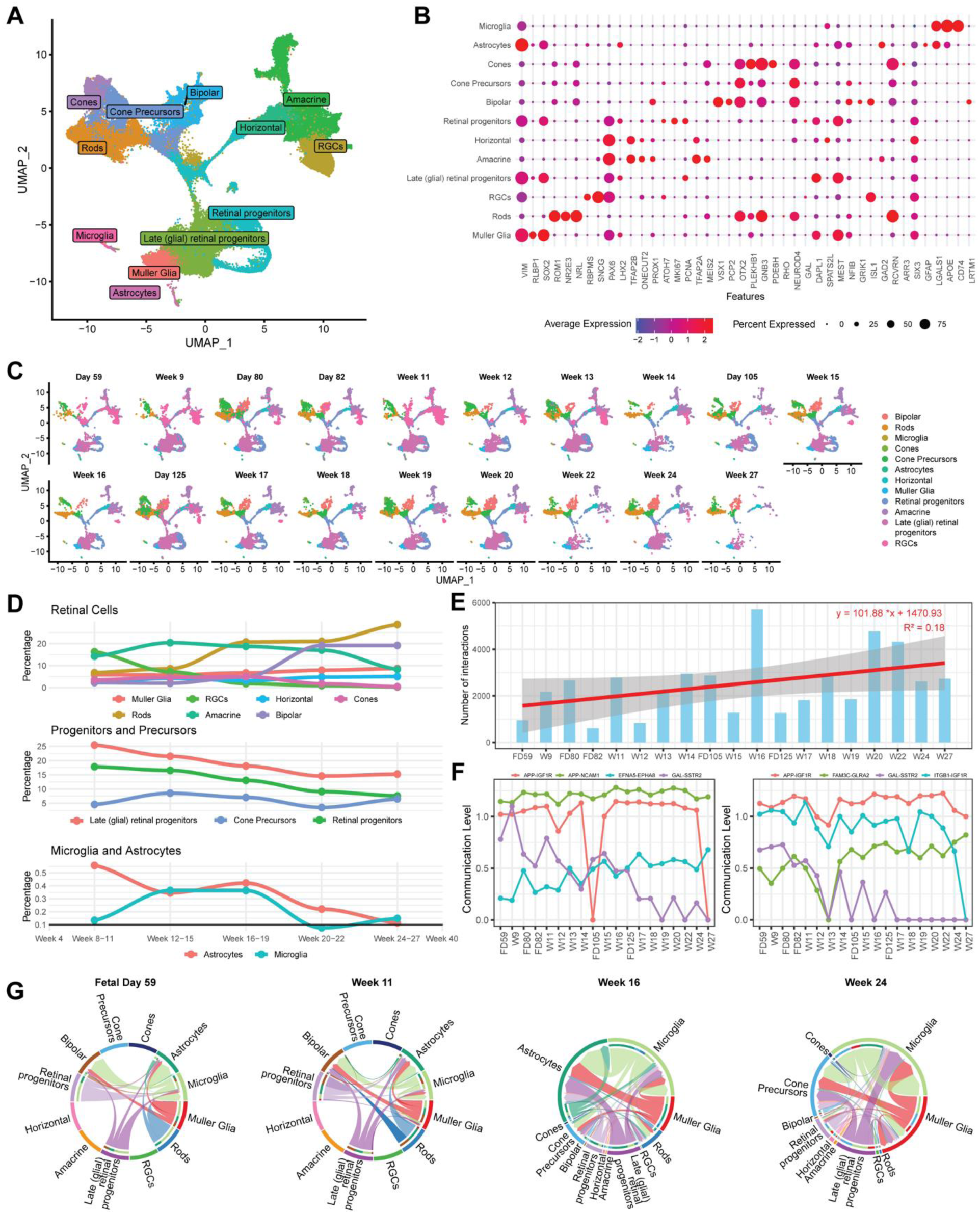
Cellular Composition and Communication in Human Fetal Retinal Development. **(A)** Integrated UMAP embedding demonstrating the cell types and their distribution in the atlas of healthy human fetal retina from fetal week 8 to 27. **(B)** Dotplot representing the cell type-specific genes correlation with the annotated cell types in the dataset. **(C)** Dimplot split by time points shows differences in cell type enrichment during the development. **(D)** Cumulative cell type frequency plots demonstrate the timeline of cell types origin and dynamics in the retina for the major cell classes (top), progenitors and precursors (middle), and microglia and astrocytes (bottom). **(E)** Bar plot demonstrating the increasing number of interactions in the retinal development with regression line (in red) and standard error (in gray). The number of interactions represents the sum of every unique ligand-receptor interaction found to be present with the CellChat cell-cell interactions analysis. **(F)** Patterns of communication dynamics for representative ligand-receptor pairs from the total interactions obtained using an LRLoop analysis for the signaling from (left) and to (right) RGC across the development. **(G)** Chord diagrams for SPP1 ligand-receptor pairs across development, generated using CellChat analysis, show patterns of increasing signal complexity (from fetal day 59 to week 11) and new signaling cell types appearing (from week 11) during development. The diagrams visualize the interactions, where the outer circle stands for the cell class, the sector size demonstrates the contribution of the cell class to the signaling, and the inner circle shows the signaling receivers for every cell class. The arrows point in the signaling direction from sender to receiver, where the thickness of the arrow codes signaling strength.

Separating these data according to their canonical timepoint (**Figure 1B**), we show different cellular compositions during each stage of retinal development, with RGC comprising the largest percentage of retinal neurons from weeks 8 to 11 (**Figure 1C-D**). As the retina further develops, the percentage of retinal progenitors and precursor cells is reduced while the other populations of retinal neurons increase (**Figure 1D**). Together, these data based on single-cell RNA sequencing match those quantifying the cells arising at different developmental stages(*29*). However, unlike the histological and cell sorting approaches used to understand neuron birth order in the retina, by establishing a human fetal retinal atlas, we can explore cellular communication to and from each individual cell population, as well as their gene profiles and the tissue as whole (*18*).

The cell-cell communication analysis shows the increased number of interactions throughout development (**Figure 1E**). We can broadly classify the strength of individual ligand-receptor pair interactions into four groups: starts high, remains high; starts high, goes lower; starts low, goes higher; and starts low, remains low (**Figure 1F**) – “high-high”, “high-low”, “low-high”, and “low-low”. We can hypothesize which interactions are important to best support a specific cell class by matching these patterns to the prevalence of that cell class (**Figure 1D**). For example, because the prevalence of RGC is greatest during early development, we predict that receptor-ligand pairs classified as “high-low” is necessary for RGC development. Lastly, we can also delve deeper into a single to study the involvement of individual cell populations (**Figure 1G**), highlighting the importance of those cell-cell interactions during development. A complete panel of all those cell-cell interactions is available in **Figure S3**.

### Retina Development Analysis Demonstrates Key Trajectory States, Bottlenecks, and Cell Classes Origin Points

Trajectory analysis (RNA Velocity(*12*), CytoTRACE2(*13*), Mellon(*14*)) shows how retinal cell classes develop from their progenitors in glial and neuronal fates, as well as what cell classes arise from which type of progenitors (**Figure 2A**), which aligns well with the current paradigm of intrinsically different retinal progenitors. (*1*), capable of producing different types of retinal neurons. The common retinal progenitor, which contains proliferating subpopulation, gives rise to two main branches: glial progenitors and neuronal progenitors. The pattern correlates with the current retina development paradigm previously shown using scRNAseq methods(*4–6*).

**Figure 2.**
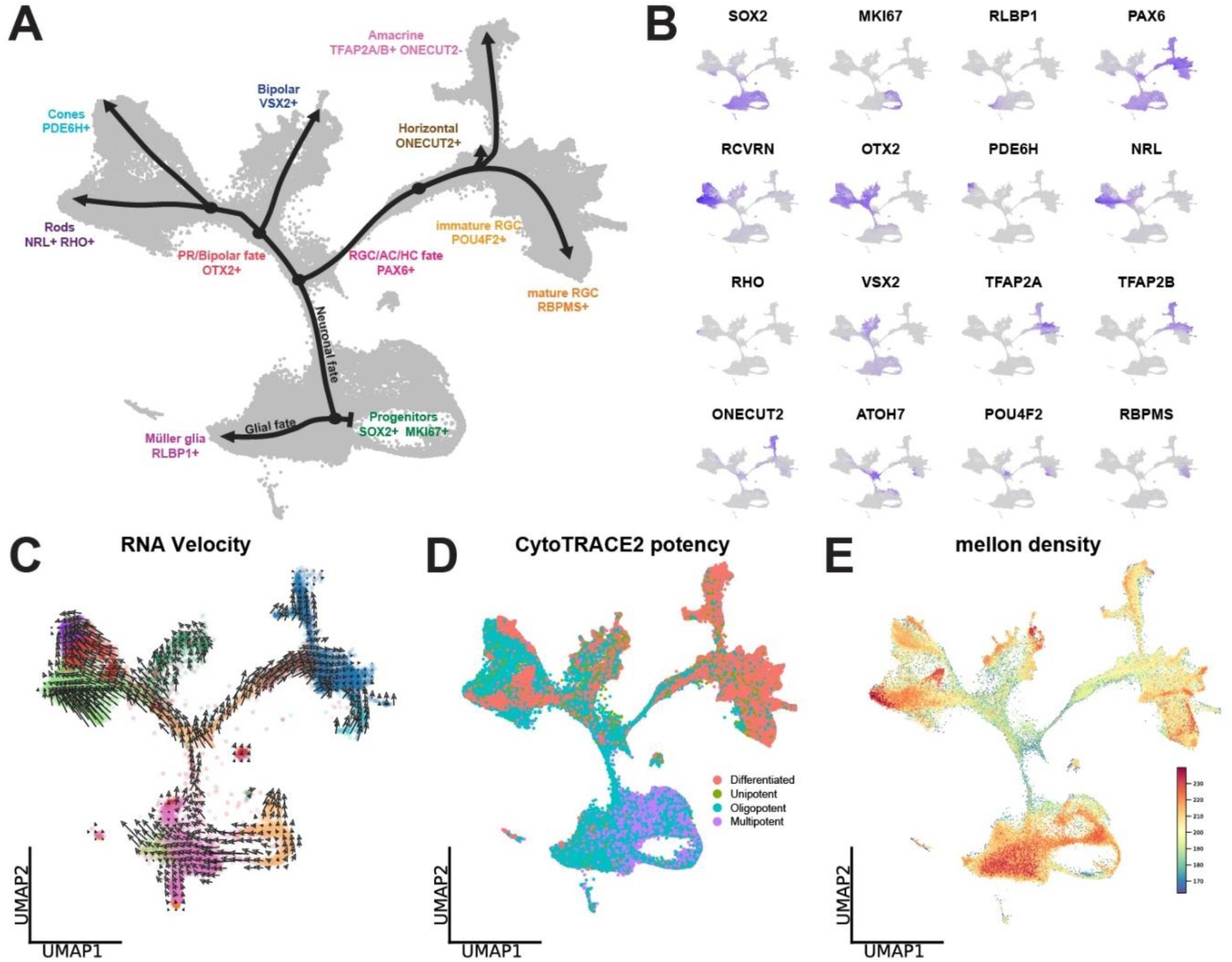
Trajectory Map of Human Fetal Retina Atlas. **(A)** Embedding demonstrates the main branches, transitory states, and terminal points of retina development present in the retina atlas with their gene profiles. **(B)** Feature plots demonstrate the genes relevant to every state on the trajectory map. **(C)** RNA Velocity demonstrates the direction of the trajectory for the retina atlas. Each arrow shows the local trajectory from progenitor to progeny based on spliced/unspliced RNA score. The arrow length codes the transition velocity, and the arrow direction points to the desired state of transition. **D)** CytoTRACE2 output results showing the potency levels for every cell in the atlas. Color codes the potency of the cell. **(E)** Mellon output showing the cell density at every embedding coordinate, demonstrating the areas of fast and low cell fate transition speed, as well as the terminal points.

The glial gives rise Müller glia (RLBP1+) and neuronal gives rise to all the retinal neuron classes. The neuronal progenitor splits into two fates: “RGC/AC/HC” and “bipolar, cones, and rods”. RGC/AC/HC fate is highly PAX6 positive, while photoreceptors/bipolar fate expresses OTX2. These two fates will give rise to the corresponding cell classes: RGC, AC, HC, and bipolar, cones, and rods. It is important to mention that for all cell classes we can identify a committed single-track “precursor” with exclusive transcriptional signature, that is acquired before full maturation. In this manner, cones are PDE6H+ yet before they become transcriptionally mature, and RGC go through POU4F2+ stage before acquiring RBPMS expression(*30*). Rod population is NRL+ RHO+, bipolars are VSX2+, HC can be defined as ONECUT2+TFAP2A,B+, and AC as ONECUT2- TFAP2A,B+ (**Figure 2B**).

Besides that, we observe two separated clusters of microglia/myeloid cells (CX3CR1+) and astrocytes (GFAP+). These classes, not being aligned and connected with the developmental trajectory and other classes, confirm microglia and astrocytes non-retinal- progenitor origin(*31, 32*).

We applied three independent strategies/packages to study cell trajectories at global and local scale to understand the general trajectory, and its local perturbations. We utilize RNA Velocity, a method of cell fate trajectory reconstruction (**Figure 2C**). This method allows us to resolve spliced/unspliced transcripts to build local trajectory arrows from the more unspliced cells to the cells with more spliced transcriptomes.

By combining this method with CytoTRACE2 and Mellon (**Figure 2D, E**), we enrich our understanding of cell fate trajectory and demonstrate the alignment of stemness/potency, density, and RNA Velocity trajectory.

The analysis at the canonical timepoints by CytoTRACE2 confirms the existing paradigm of retinal neuron maturation with RGC/AC/HC fate mostly complete by week 27 and “photoreceptor/bipolar” fate still undergoing maturation (**Figure 2D**).

Cell density analysis by Mellon shows the cells to be highly concentrated at the initial and terminal states of the development (**Figure 2E**) and low density at the transitory states, suggesting lack of obvious bottlenecks during development. This can be explained by mutually non-exclusive hypotheses: 1) the cells born at any time point do not have to wait for the later time point or complete cell class to become mature - cells are intrinsically sufficient – development is pre-programmed and self-supported, 2) the variety of cell maturation states can be sufficient to support the maturation of the immature cells - extrinsic auto- and paracrine signals are redundant and can come from any of the existing cell classes. In any scenario, the trajectory, potency, and density findings show that the retinal developmental system is more of an independent closed system, where maturation is self-guided by both cell-intrinsic and cell-extrinsic mechanisms.

### Pseudotime Analysis Reveals Developmental Heterogeneity and Key Maturation Stages of Retinal Ganglion Cells

The developing neural retina presents a unique system that contains developmentally heterogenous cell classes at each canonical timepoint(*33*). We explored this spectrum using RGC development trajectory as a substrate. To disentangle the maturation state diversity at each canonical time point we performed a pseudotime analysis on the ForceAtlas2 embedding using the scFates package cell fate trajectory reconstruction pipeline(*11*). It aligns the cells from the least to the most mature (**Figure 3A, Figure S4**) to the single-track developmental path. As expected, we observe that each canonical timepoint contains RGC at various developmental stages (**Figure S4**) and the RGC specification and maturation is continuous with variable gene expression along the pseudotime trajectory from RGC precursor to the most mature RGC cell (**Figure S5**). We then separated the continuous pseudotime timeline into the sectors from t_1_ to t_7_ for further analysis following the Leiden clustering algorithm(*34*) (**Figure 3B**).

**Figure 3.**
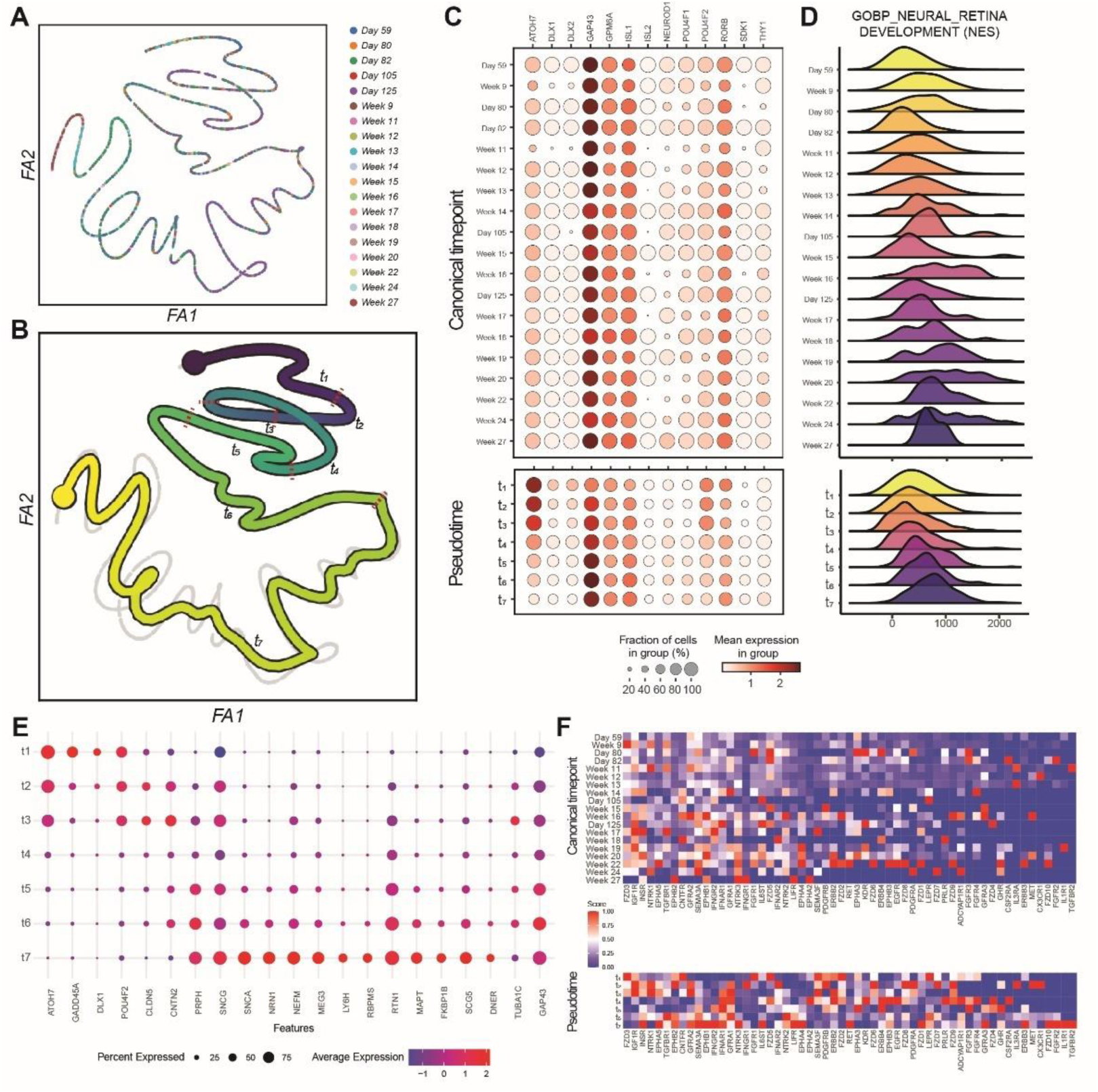
Temporal Gene Expression Dynamics of Retinal Ganglion Cell Differentiation During Human Fetal Development. **(A)** ForceAtlas2–based embedding of retinal ganglion cell from the fetal retina atlas. **(B)** Cell fate trajectory reconstructed with the scFates approach, with pseudotime values in color. For further analysis, pseudotime values were separated into seven sectors based on the automated predictive approach. **(C)** Dotplot for the genes involved in the neural retina development shows the expression differences and gene gradients between the two approaches: the canonical time points separation on the top and pseudotime separation (that may include multiple canonical time points in every pseudotime sector) on the bottom. **(D)** Ridge enrichment plots for the neural retina development pathway upon GSEA analysis demonstrate the difference between the canonical timepoint- and pseudotime-based approaches. **(E)** Dotplot showing the expression profiles of pseudotime states of RGC. Color codes the scaled and normalized expression values, with blue showing the lowest and red color – the highest values. **(F)** Heatmap showing the expression patterns of neurotrophic receptors across the canonical time points and pseudotime sectors. Color codes the scaled and normalized expression values, with blue showing the lowest and red color – the highest values.

To confirm that pseudotime ordering allows better maturation resolution, we compare the canonical and pseudotime expression profiles for select genes from the neural retina development pathway (GOBP_NEURAL_RETINA_DEVELOPMENT). We demonstrate that select genes, such as ATOH7, GAP43, and POU4F2, show a random or stable expression pattern if grouped by canonical timepoint but a gradient in their expression if split by pseudotime sectors (**Figure 3C**). Using Gene Set Enrichment Analysis, we visualized the complete neural retina development pathway, containing 80 genes, and demonstrated the same switch for non-paternal to gradient expression during development (**Figure 3D**). Similar gradient expression patterns can also be observed in several tissue- and cell-level RGC pathways during development (**Figure S6**).

We also employ the pseudotime and canonical time to disentangle cell intrinsic and cell extrinsic factors in development: the gene expression at canonical time point defines tissue microenvironment (extrinsic factors), and gene expression in pseudotime determines inherent cell development force (intrinsic factors).

We describe the pseudotime states of RGC that correlate with their maturation status (t_1_ – least mature, t_7_ – most mature) and demonstrate the unique transcriptomic profile of each pseudotime stage (**Figure 3E**). The least mature RGC_t1_ are characterized by ATOH7, GADD45A, DLX1, and POU4F2 expression, while the most mature RGC_t7_ cells express GAP43, TUBA1C, DNER, SCG5, and LY6H. We observe the decreasing (ATOH7, POU4F2) and increasing (NEFM, MEG3, RTN1) gradients of gene expression with RGC maturation. We identified several de novo genes previously not correlated with human RGC development: DNER(*35*), DLX1(*36*), CLDN5(*37*), and CNTN2 that were previously known to play a role in murine retina development but not in human. MEG3, the TGF -β regulator(*38*), and MAPT, known to be the microtubule stabilizer, related with early neuronal disfunction(*39*), are the new genes participating in human RGC development. FKBP1B(*40, 41*), SCG5(*5*), and GADD45A(*9, 42*) were described as human retina but not RGC development-related genes. RTN1 was known to play a role in RGC development(*43*), while NRN1 role was known to be survival-oriented(*44*), and PRPH was described as ipRGC marker(*45*).

To predict soluble factors necessary for each development stage, we built a heatmap of all known neurotrophic receptors with pseudotime and canonical time staging (**Figure 3F**).

The expression of receptors on RGC did not show any recognizable patterns at canonical timepoint, probably due to developmental heterogeneity of RGC. Pseudotime analysis allowed to identify several trends with high expression early in maturation and low at the end stages and vice versa. For instance, presence of FZD3 and IGF1R on immature, but not on mature RGC (*46, 47*), becomes apparent only in pseudotime with their highest expression in the t_1_-t_3_ state (**Figure 3E**). We also demonstrate CNTFR and SEMA3F receptors to be expressed on the surface of immature RGC, giving the insight into RGC role as a receiver of WNT, IGF, CNTF, and SEMA3 signaling in the developing retina. The most mature RGC_t7_ tend to carry more inflammatory profile by expressing LIFR, IL6ST, IL1R1, and TGFBR2.

### Whole Transcriptome Pseudotime Ordered RGC Analysis Demonstrates the Key Developmental Drivers

The maturation of RGC is a continuous and dynamic process involving a broad spectrum of genes and signaling pathways(*29*). To further investigate and characterize the maturation of RGC, we switched from studying currently known genes and pathways to the whole transcriptome analysis (**Figure 4A**). On the subset of RGC, ordered by canonical timepoints (Week 8 to Week 27) and pseudotime stages (t_1_ to t_7_), we show the peak timepoint for every gene, where its expression is the highest between canonical timepoints (**Figure 4B, left**), and pseudotime stages (**Figure 4B, right**). Particular peaks, like Day 105 or Week 22, are most likely related to batch effect, which complicates studying canonical timepoints.

**Figure 4.**
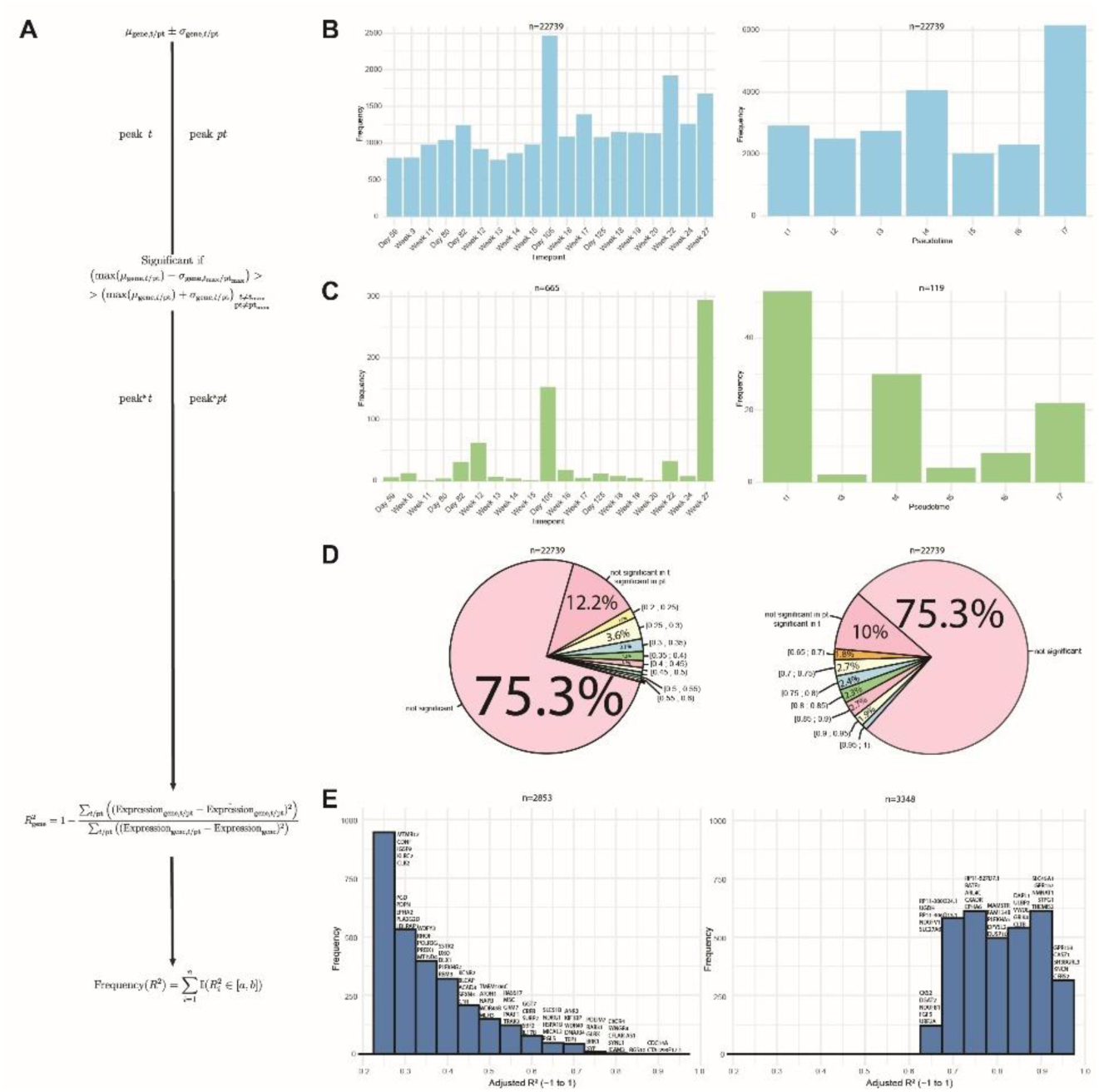
Unraveling Cell States of RGC with Pseudotime Approach Results in Enhanced State Profiling. **(A)** The approach generated and applied to filter out the genes important for canonical timepoint and pseudotime staging of RGC during their development. **(B)** Barplots demonstrating the distribution of the RGC total genes peak expression between the canonical timepoints (left) and pseudotime stages (right). **(C)** Barplots demonstrating the distribution of the RGC significant genes peak expression between the canonical timepoints (left) and pseudotime stages (right). **(D)** Pie charts showing the distribution of regression values for the total genes in RGC between canonical timepoints (left) and pseudotime stages (right). **(E)** Barplots demonstrating the significant genes (top 5 per bin) and their regression values between the canonical timepoints (left) and pseudotime stages (right) for RGC.

Upon adding the dimension of significance and filtering out the whole transcriptome to significant genes, we see a different pattern for canonical time points and pseudotime stages (**Figure 4C**). For the canonical time points, the RGC at Day 105 and Week 27 express the highest number of genes. Yet, no information can be accessed on RGC maturation states since canonical time points resemble a mix of maturation states of RGC, as shown in **Figure S4**. On the other hand, we see the most significant genes concentrated in the t_1_, t_4_, and t_7_ pseudotime stages of RGC, the least, the mid, and the most mature states. Such a distribution demonstrates the most gene-enriched maturation stages.

We would like to understand the peak timepoint or pseudotime stage for each gene and what genes are essential for development. Following this idea, such a gene would either have a pattern of increasing or decreasing expression with development. Since canonical time points represent a complex structure of maturation states mixture, we suppose that pseudo time ordering may unravel this heterogeneity by ordering cells from their canonical time points into maturation state. We performed regression analysis on the whole transcriptome to identify the maturation-related genes (**Figure 4D**). Most genes are insignificant for canonical time points (87.5%) and pseudotime stages (85.3%). However, the regression values for the canonical time points vary from 0.2 to 0.9, with the dominant part in the diapason of 0.2 to 0.6, with the total number of significant genes being 2853. (**Figure 4E**). When performing regression analysis using pseudotime stages order, we demonstrate that most significant genes (n = 3348) fall in the window of 0.65 to 1 regression values. For each bin (step = 0.05), we present the top 5 significant genes, with the most critical genes with the highest regression values to be CDC14A, CTA293-F17.1,

RGS10, CXCR4, SYNGR4 for canonical time points, and GPR153, CASZ1, SH3BGRL3, KNCN, CERS2 for pseudotime stages. Some of the genes described upon pseudotime stages regression analysis were previously known, while some with the strongest maturation correlation have no evidence from the literature(*48–50*).

Aligning RGC in pseudotime also allows the filtering out of potential batch-effect-related artifacts. For instance, false positive conclusions on the major timepoints of RGC development to be Day 105 and Week 22 could be described as sequencing coverage or sample composition issues. These are resolved upon pseudotime alignment by 1) switching from sample and replicates structure into continuous trajectory, 2) averaging the noise from sequencing depth and coverage into large pseudotime stages of RGC maturation.

### RGC contribution to the retinal microenvironment changes with their developmental stage

We explored RGC population as a donor of extracellular signaling molecules, relevant for retinal development and maintenance. During the RGC maturation, new cell classes start arising, making the retina system more and more complex. RGC also require autocrine regulation for their maturation(*51, 52*).

We performed a CellChat analysis of every canonical timepoint (**Figure S3**) and identified the top signals that RGC send to other retinal cell classes (**Figure 5A**). We also separated the contribution of pseudotime RGC stages at each canonical timepoint and identified the major signaling donor (**Figure 5B**). For example, we demonstrate that the most frequent recipients of RGC signaling are RGC themselves. Also, for the cell classes such as bipolar, cones, and amacrine, the prevalent RGC state sending the signal is RGC_t7_, the most mature state of RGC. RGC of all developmental stages also communicate with retinal progenitors by sending ncWNT, NOTCH, SEMA3, and EPHB signals (**Figure 5A, B**). We observe multiple signaling pathways to be sent by RGC to other cell types throughout the RGC maturation: astrocytes get the ADGRG and ADGRE support from t_1_ to t_7_ states of RGC, cones receive GABA-B from t_3_ to t_5_, microglia start receiving prostaglandin signaling from t_3_ to t_7_ (**Figure 5C**).

**Figure 5.**
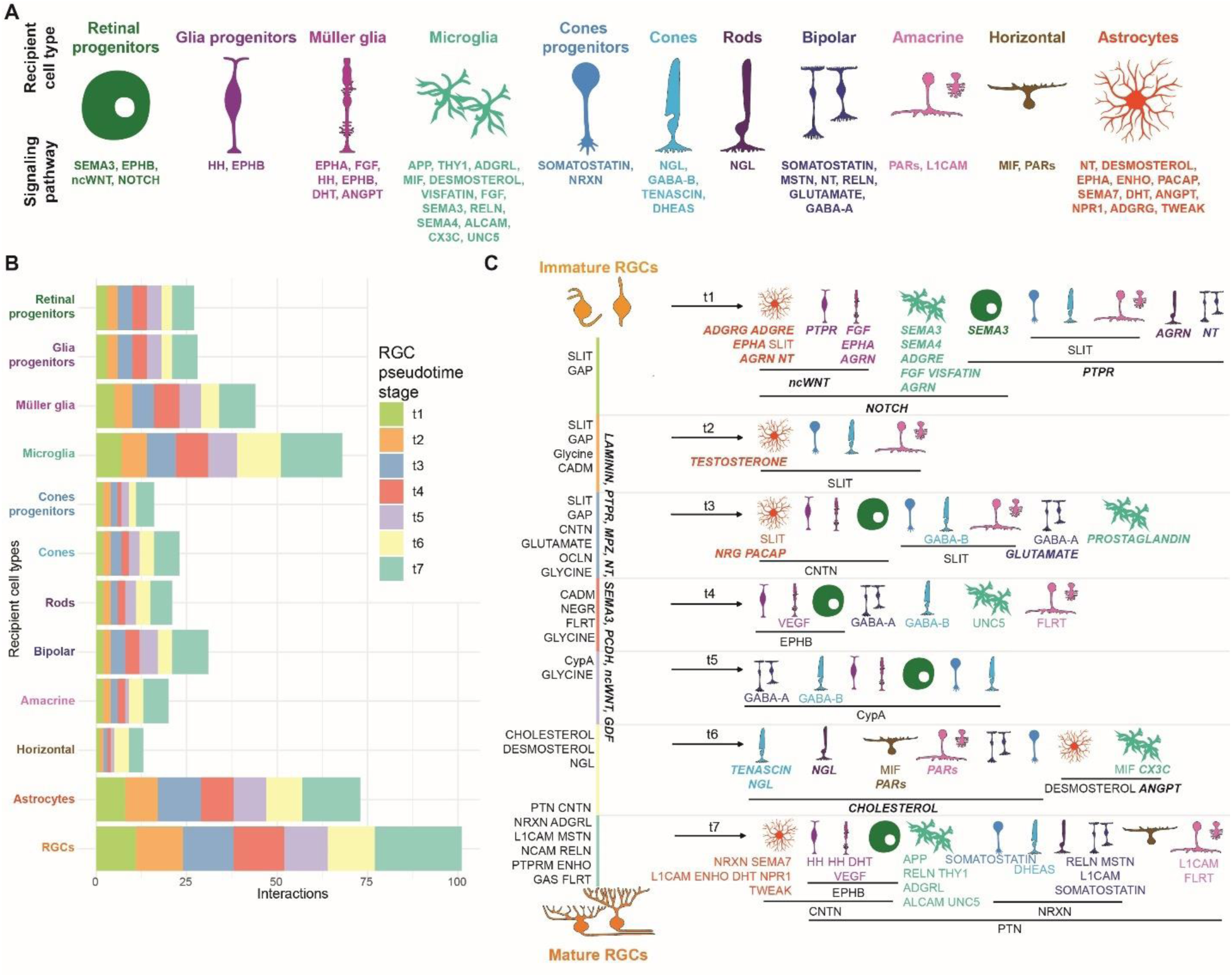
Cell Extrinsic Dimension of RGC as Signaling Donor in Pseudotime Staging Resolution Demonstrates the Dynamical Interactions Network with Retinal Cell Classes. **(A)** The major signaling pathways sent from RGC to different cell classes present in the fetal retina atlas. **(B)** Barplot demonstrating the distribution of RGC pseudotime stages sending the signals to retinal cell classes. **(C)** Scheme showing different RGC pseudotime stages and the signals they send to different retinal cell classes. Signaling pathways described on the left from the vertical line from immature to mature RGC are the autocrine regulatory pathways. Signaling on the right from the vertical line demonstrates the signals RGC sends in the resolution of pseudotime staging from t_1_ to t_7_. Signaling shown in bold italic font is present in further pseudotime stages upon its first appearance. Signaling in regular font is sent from the relevant RGC pseudotime stage only.

RGC autoregulate at the early stages (t_1_-t_3_) of development by SLIT and GAP, at the mid stages (t_3_-t_5_) by glycine, and at the late stage (t_7_) by CNTN, NRXN, and NCAM production. RGC also support themselves throughout the whole maturation by LAMININ, PTPR, MPZ, NT, SEMA3, PCDH, ncWNT, and GDF (**Figure 5C**).

### Human fetal retina atlas as a reference tool for automated data annotation

We have downloaded and reanalyzed the day 45 human induced pluripotent stem cell (IPSC) -derived retinal organoids data(*6, 50*). The resulting object was annotated by three approaches: 1) manual annotation using known marker genes; 2) automated annotation using a new fetal retinal atlas; 3) automated annotation using an adult retinal atlas **(Figure S7A)**.

Upon manual data annotation, we achieved the resolution of progenitor, transitory, amacrine, immature and mature retinal ganglion cells **(Figure S7B)**. We further used Azimuth to perform automated data annotation of the sequencing data using human fetal retina atlas that resulted into higher cell class resolution compared to the manual approach **(Figure S7B)**. We identified common retinal progenitor, glial progenitor, astrocyte, horizontal, amacrine, and retinal ganglion cells.

Finally, we have downloaded and reprocessed the dataset on human adult retina to use it as a reference for the annotation described in our previous paper(*50, 53*). The resolution achieved included the majority of cells to be annotated as rod photoreceptor cells, with the rest (<15% from the total number) to be identified as astrocytes, Müller glia, microglia, amacrine, and horizontal cells **(Figure S7B)**.

Compared to the known retina markers **(Figure S7C)**, we noticed that the annotation performed using human adult retina does not correlate with the markers expression: for instance, the majority of the cells are annotated as rods, while the cells express SOX2, MKI67, ATOH7, POU4F2, TFAP2A,B. These markers mostly represent retinal progenitors, RGC precursor, and RGC.

We further compared the annotation approaches **(Figure S7D)** and found manual and automated annotation using human fetal retina atlas to overlap in the identities and frequencies of cells in each identity compared to the automated annotation using human adult retina. Both manual and automated annotation using human fetal retina of these identified amacrine, photoreceptor, progenitor, and RGC classes correctly. The automated fetal atlas annotation also identified astrocyte, cone precursor, and horizontal cells, while we could not resolve them with manual annotation.

On the other hand, the automated annotation using human adult retina identified such cell classes as microglia and Müller glia cells, that are not supposed to be present in the day 45 hIPSC-derived retinal organoid(*6*).

## Discussion

In this study, we focus on human retina development and the methods to unravel the heterogeneity of retinal cell classes’ maturation states at any given timepoint throughout development. Multiple anatomical, electron microscopy, evolutionary, molecular, and scRNAseq studies(*4–10, 54–59*) demonstrate human retina development map with its transitory states, bifurcations, and differentiated cell classes. We confirm the current concept of retinal development and support it by bringing additional dimensions to the understanding of intrinsic and extrinsic parts of the maturation function.

Our computational studies validate the current paradigm of intrinsically different retinal progenitors (*1*) in an independent, unsupervised manner. In human developing retina, SOX2+ progenitor population separates to glial and neuronal progenitors, forming two separated fates. Our analysis highlights a list of major cell classes, including the unique ones for the development condition (progenitors, transitory states), and their exclusive markers allowing easier targeted downstream analysis of them in both ‘dry’ and ‘wet’ biology.

Compared to the previous studies(*6*), we demonstrate that it is possible to identify and analyze the immature RGC subpopulation in the dataset that is negative for “classic” early RGC markers: GAP43, SNCG, POU4F2, or NEFL. On the other hand, RBPMS, being a marker of mature RGC, cannot be used as a universal RGC marker when studying retinal and RGC development(*10*). We also show the canonical RGC-related genes behavior during RGC maturation, as well as demonstrate the profiles for RGC at different stages of maturation. We demonstrate that immature RGC can be observed in the human retina after fetal week 13(*4*).

The human fetal retinal atlas established herein can be used not only in the discovery of novel factors but also for targets to affect any number of cellular processes, including cell migration, maturation, and synapse formation. This search may be started by studying the dynamics of cell class development and understanding the cell fate drivers of timepoint-specific factors. The current version of the atlas offers a broad spectrum of high- throughput data in the aspects of cell-cell interactions, regulons, and transcriptional motifs for every cell class at every time point. With that, many potential applications exist that increase the borders of factors search from intrinsic gene expression changes to ligand- receptor pairs, communication patterns between cell classes, variable regulons, and further motifs analysis in connection with other available methods, such as ChIP-, ATAC- seq, GWAS, and Hi-C. Since this atlas demonstrates the first evidence of regulons and motifs (list available upon request) in human developing retina, it could give clues for population-related studies.

What we found is that the novel factors search may be performed without the connection with canonical time points. Some cell classes, like RGC, are born over a specific period of time in a continuous fashion. As a result, at any particular moment, the RGC population within a human retina contains a range of developmental stages and demonstrates the gene dynamics that could not be seen using canonical time points yet could be observed using pseudotime methods. A significant part of our work is devoted to the idea of unraveling the maturation states heterogeneity. It is essential to understand the development in its maturation complexity and recapitulating its variety and heterogeneity is the approach that looks to be the closest to the native model.

With that application, utilizing the methods that could unravel developmental heterogeneity, such as pseudotime, potency, or other development-related function, could help us uniform the multidimensional heterogeneity. We show that the application of pseudotime analysis methods and further cell reordering based on it leads to significant changes of the analysis results compared to canonical timepoints analysis. Pseudotime as a grouping parameter instead of time points unravels the state heterogeneity and allows a direct search of state-related targets. This may be applied not only for the default differentially expressed genes analysis but to the same methods of ligand-receptor pairs, regulons, and motifs analyses. When canonical timepoints demonstrate the mixture of cell states, a lot of noise comes into understanding the development-related parameters. Arranging the cells by their maturation state into an artificial system helps to understand the development drivers and to profile the development intrinsically and extrinsically. Besides that, the application of pseudotime allows the switch from classical discrete clustering, usually applied in bioinformatics, to the maturation state continuity. Our choice of pseudotime stages (from t_1_ to t_7_) is used to visualize and compare the differences between pseudotime and canonical timepoint approaches. Yet, the pseudotemporal resolution could be increased to as many stages as relevant, or performed in continuous manner, while it worth mentioning that to increase temporal resolution in canonical timepoints new biological samples have to be collected.

We also observe that within the batch effect correction, it is impossible to adjust for unique sample perturbations. As in our case, some of the timepoints demonstrate significantly more genes to be dominant across all the timepoints (Day 105, Week 22), it is mostly due to the fundamental issue of the batch effect underlying not in the expression differences directly but the sample composition variation. Application of pseudotime here allows to switch from canonical timepoints, each representing a unique sample, into the computational model of continuous pseudotime maturation course, not dependent on the sample variation anymore but the maturation states variation covered in the atlas.

We demonstrate these results for RGC, yet the rest of the cell class could be analyzed following the same pipeline. Further cell fate trajectory reconstruction and branch early- and late-driving genes analyses using RNA Velocity or scFates may be applied to increase the resolution from cell class to cell type and identify the type-related drivers, especially for amacrine, bipolar, and RGC, highly heterogeneous in the cell types. For one to perform such a study, it is crucial to include the progenitors or precursors (like ATOH7+ cells within RGC trajectory) of the cell type of interest in the analysis to have a broader spectrum of maturation. To identify the progenitors or precursors within the atlas for the specific cell class, the same methods for cell fate trajectory could be applied (RNA Velocity, CytoTRACE2, Mellon, scFates). We also provide the exclusive markers defining the fates to simplify this task.

This idea can be brought to its maxima by generating and correlating the maturation state for each cell class with their further rebuilding into the artificial system of the least mature, more mature, and the most mature cell from every cell class to synchronize inter-classes maturation.

The main limitation of the pseudotime analysis is the heterogeneity inherited in the biological system studied. Additional datasets replicating the same timepoints or broadening the time points window with high sequencing depth could be applied to reintegrate the atlas. The ultimate task here is in covering the most of the natural-limited developmental cell states to avoid potential false results coming from the lack of the sequencing data.

Arranging the natural-limited developmental cell states can also be performed using the other development-oriented functions. This may be the output of CytoTRACE2, RNA Velocity latent time, Monocle 2/3 pseudotime, or other ordering parameter. If applying these, the expected result may overlap between the methods, demonstrating the most conserved genes. However, each method may produce a unique, non-overlapping group of genes that requires careful evaluation and testing. For instance, switching from pseudotime to CytoTRACE2 potency output, may result in reordering the cells with the focus on potency. Even if potency has direct connection with maturation, for the non-uniform directions (neuronal progenitor to M-cone, ATOH7+ cell to ipRGC) with possible local perturbations or additional bifurcations on the trajectory, the result may differ. Switching from RNA Velocity pseudotime to latent time or vice versa may impact in temporal reconstruction of the dataset, due to different computational algorithms behind. Further adjustment of cell order may be achieved within one temporal/developmental function by changing such parameters as number of diffusion components, neighboring graph, embedding, or dimensionality reduction approach.

The atlas in its current version may be a significant resource for basic research and translational applications. The loss of RGC is one of the main causes leading to vision decline and blindness. While many strategies for vision rescue follow the idea of establishing the artificial retina model (organoids, cell culture, organ-on-a-chip)(*6, 50, 60–71*), some of them look in the direction of recapitulating the development per se. This strategy means pushing the artificial model into the same intrinsic and extrinsic state with dynamical environment shifts upon the model maturation. Therefore, the findings described provide multiple insights into fetal retina composition, cell classes origin, their trajectory that needs to be recapitulated. With the focus on RGC, we show what states the cells have to pass during their maturation.

These states, described from both cell-intrinsic and cell-extrinsic prospectives, are the native trajectory states RGC need to follow in order to become functional. A potential application of the results here can be the modulation or support of the artificial in vitro system by iterative correlation of the system to native retina state and further small molecules, ligands addition, or genes activity modulation.

Multiple papers demonstrate the correlation of the artificial system, such as organoids, with the fetal retina, using IHC, bulk RNA-seq(*33, 58, 70*). However, single-cell RNA-seq has more capacity to compare the natural and artificial systems with advanced precision and multiple dimensions of data to analyze. We demonstrate the application of our atlas as a tool for automated sequencing data annotation in comparison to manual annotation and automated annotation using human adult retina data. The master human fetal retina atlas object is easily convertible to the reference atlas format using the Azimuth package (tutorial is provided on the GitHub page) for further integration of the model sequencing data into the atlas. This will result in multiple metrics, such as automated cell identity labeling, confidence of the labeling, and similarity to the native atlas cell at the same coordinate on the embedding. Such an approach may significantly benefit in solving the common problem of cell-class-like identity of the model cells, where it becomes challenging to correlate the genetic profile of the model cell with the native fetal retina cell. If one would select a manual annotation method, our discoveries may serve as a better signature of RGC maturation states, branches and other retinal classes exclusive markers necessary for the manual cell class/type/state profiling.

It may further deepen the understanding of the model cell structure thanks to the unified embedding with the atlas. In this manner, cells from the model system will be embedded on the same coordinates as the cells are in the human fetal retina atlas, if the model system cell is similar enough to the atlas cell. Thus, it may give a hint into understanding the current model maturation bottleneck, cell class and state enrichment. Further correlation and bottleneck analysis may give a hint into potential modulation to apply within the system in order to push its maturation further. Such an iterative approach on ‘model system sequencing – reference integration – automated correlation – model system modulation’ serves a potential strategy in improving the differentiation and transplantation outcomes, since immature cells do not completer their development upon transplanting into the adult retina environment due to the lack of the resources needed for their maturation.

We also describe the complete map of cell-cell interactions within the developing human retina with the focus on RGC as both donor and recipient. It may serve the purpose to improve cell transplantation outcome by modulating the host retinal microenvironment upon treatment to form the pro-regenerative environment, resembling the developing retina(*72*). Depending on the model or state of the disease, the retinal microenvironment may need to be altered to support donor RGC migration, growth, maturation, axon extension, and increased metabolic demands. We believe further exploration of our fetal retinal atlas will lead to discovering and testing various cues to promote each of these key donor RGC functions. Nevertheless, integrating neuroprotective strategies with cell replacement therapies will likely be a pivotal direction for research and clinical applications. Lastly our work may be used to select novel combinations of pro-survival factors and test them as for RGC neuroprotection. Altogether, these findings provide promising insights into potential strategies for restoring vision lost due to RGC damage or loss in optic neuropathies, allowing us to switch from a reductionist to a holistic approach to testing the experimental hypotheses.

## Author contributions

E.K., P.B. performed the research; E.K., E.L. analyzed the data and created the figures; E.K., J.R.S. wrote the paper; and P.B. supervised the research.

---

**Figure S1.**
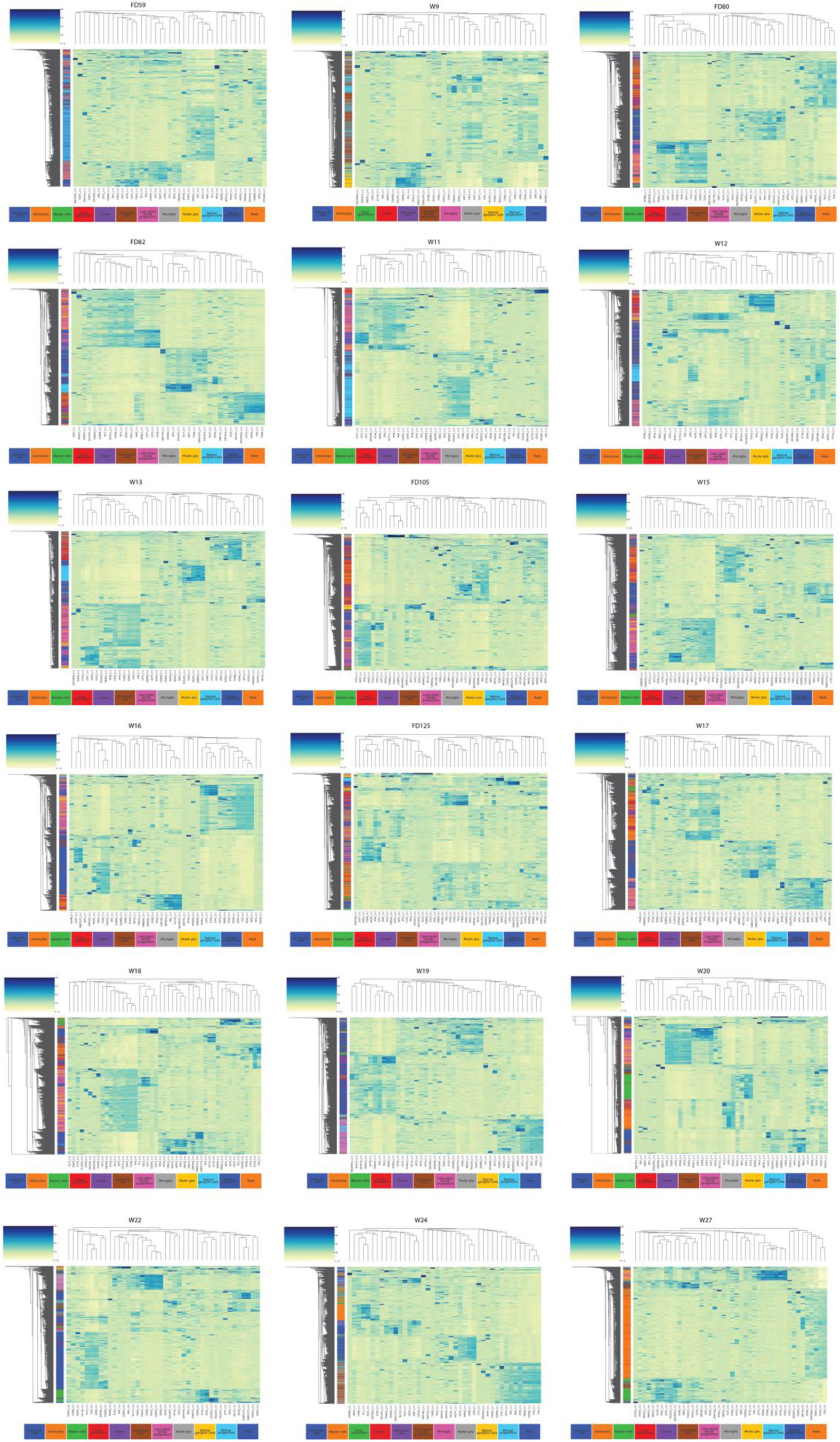
Timepoint/cell class resolution panels representing expression of variable regulons within the atlas.

**Figure S2.**
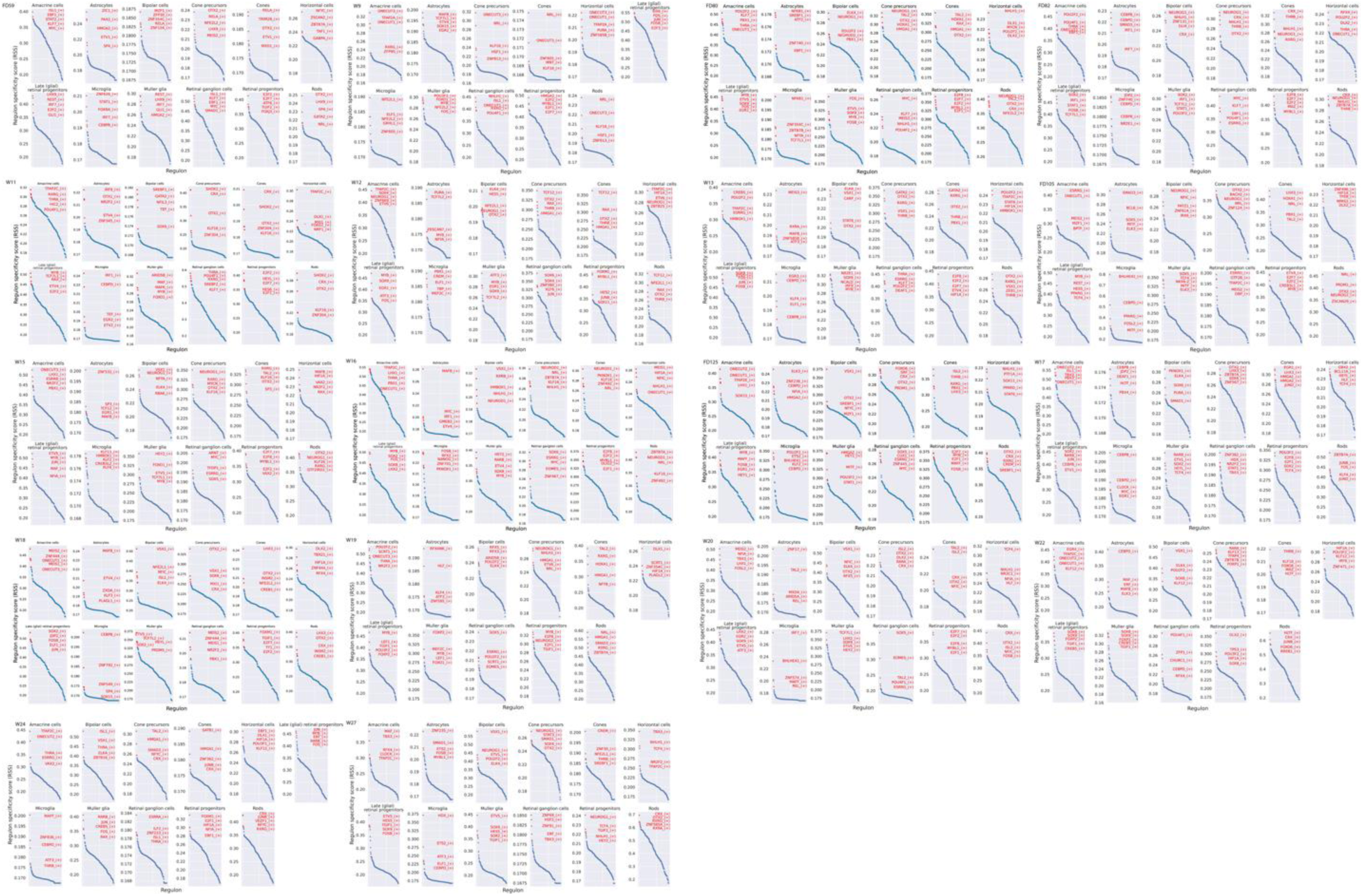
Top-5 regulons for every timepoint and cell class upon pySCENIC analysis demonstrate both conservative and variable regulons in developmental dynamics.

**Figure S3.**
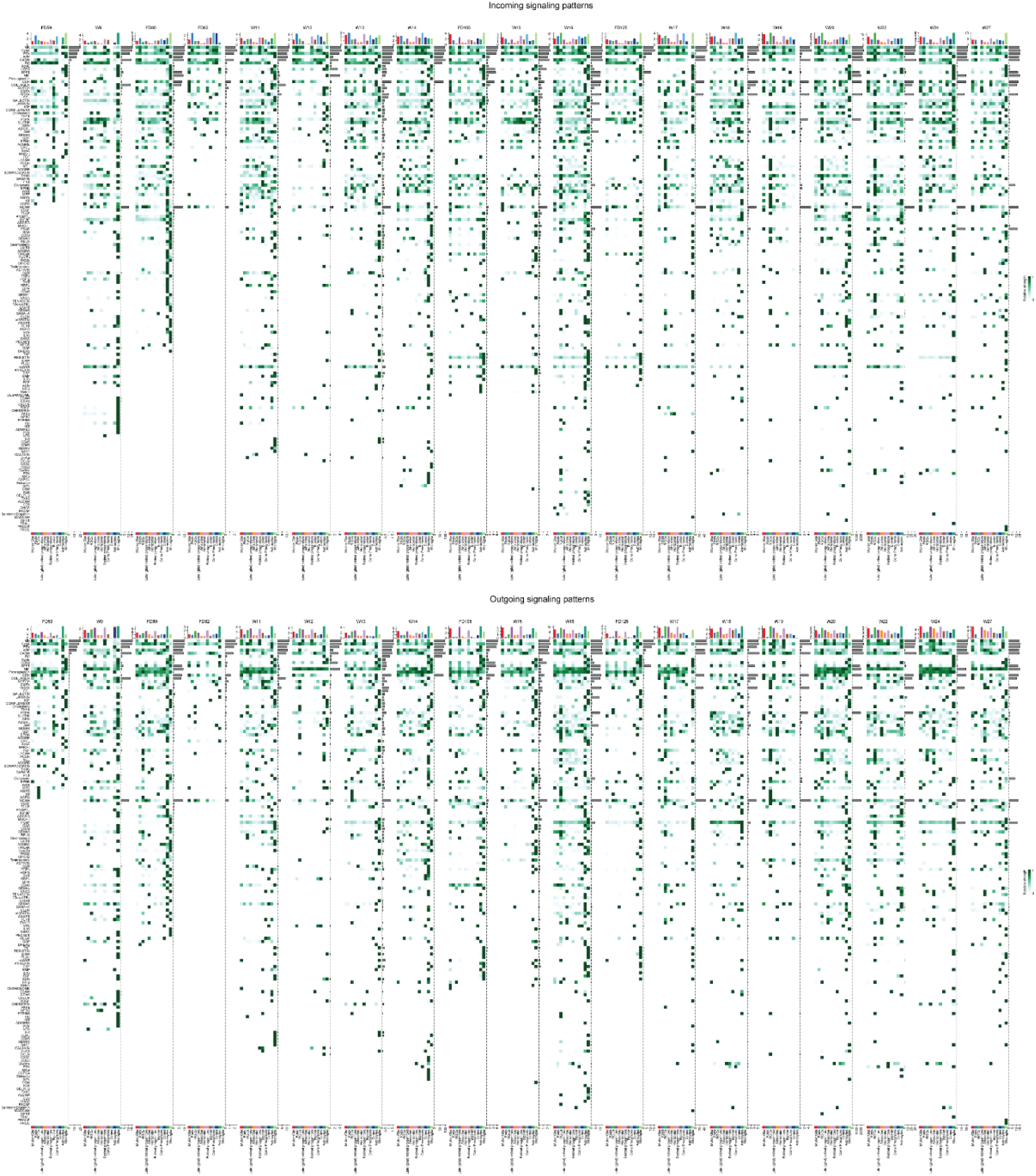
Cell-cell interactions in retinal development. Complete panel demonstrating all the incoming and outgoing cell-cell interactions for every cell class and every timepoint in the generated human fetal retinal atlas using the CellChat analysis.

**Figure S4.**
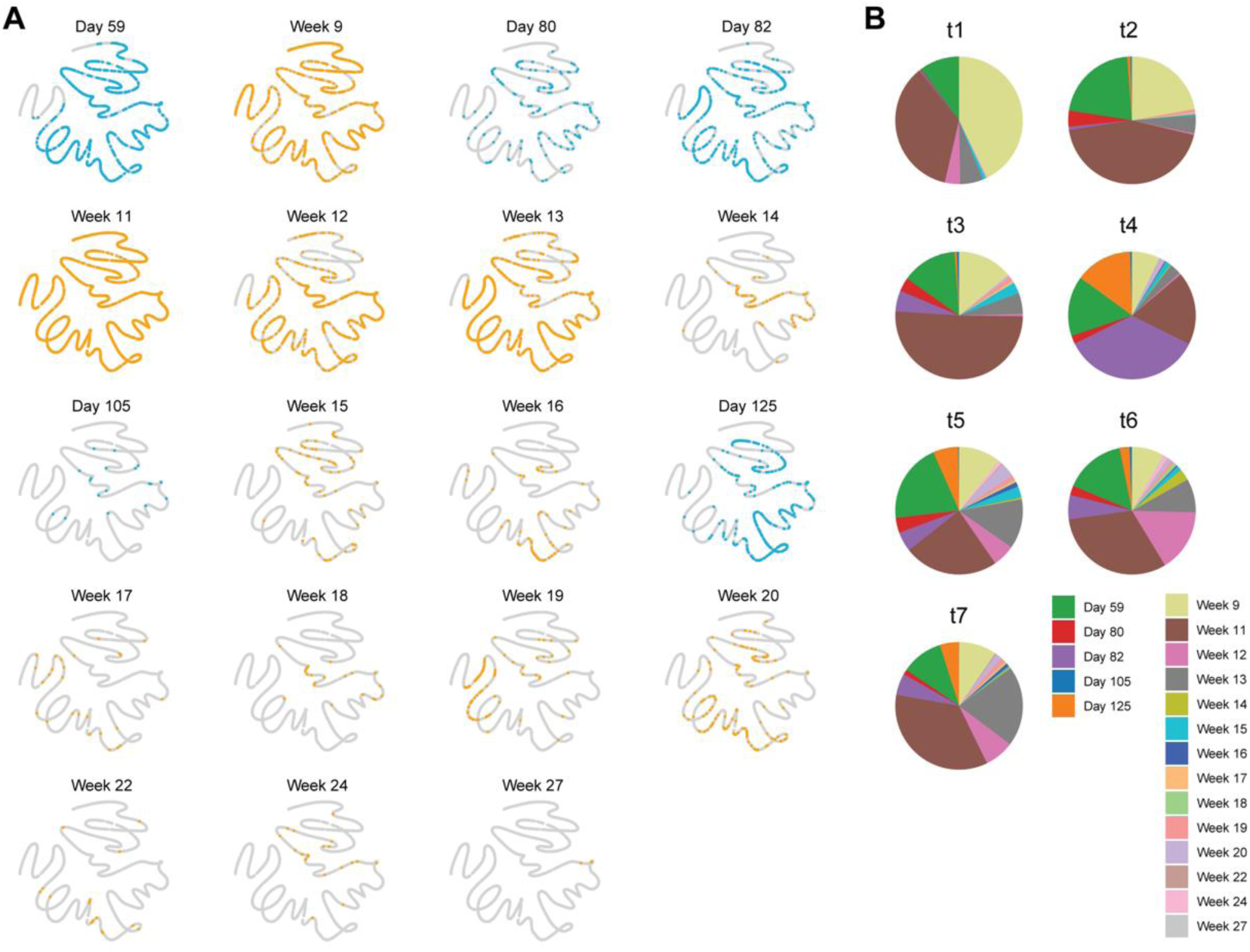
Sourcing of RGC for ForceAtlas2-based embeddings. **(A)** Distribution of retinal ganglion cells across the time points on the UMAP embedding. Color codes the public source of the data. **(B)** Pie charts showing the distribution of the time points for every pseudotime sector.

**Figure S5.**
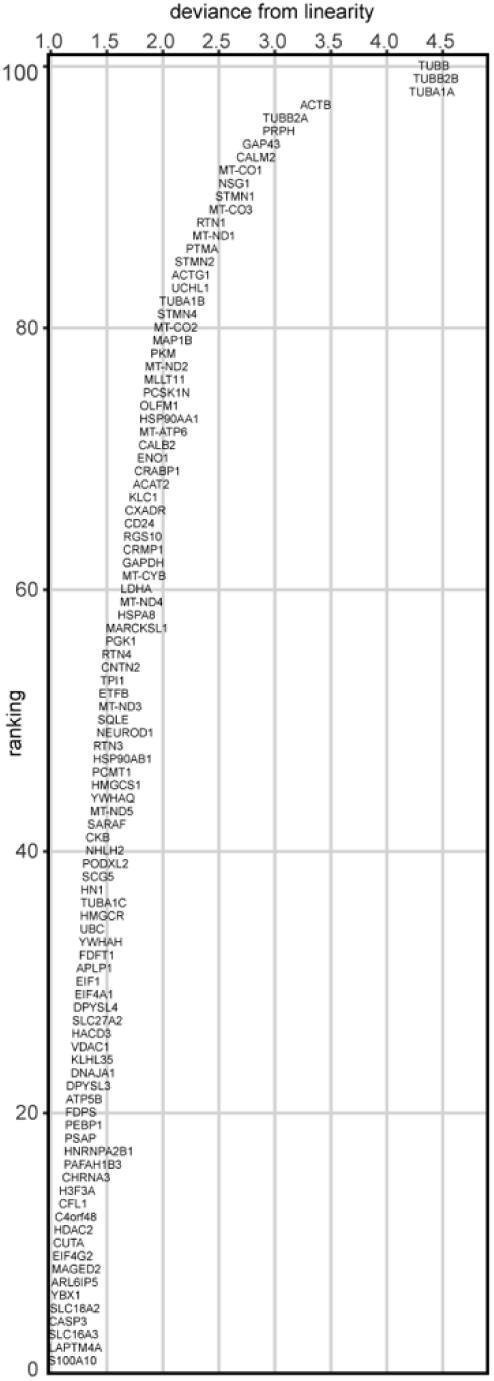
RGC variable gene expression pattern along the cell fate trajectory. The deviance from the linearity parameter codes the distance from the root to the end of the trajectory.

**Figure S6.**
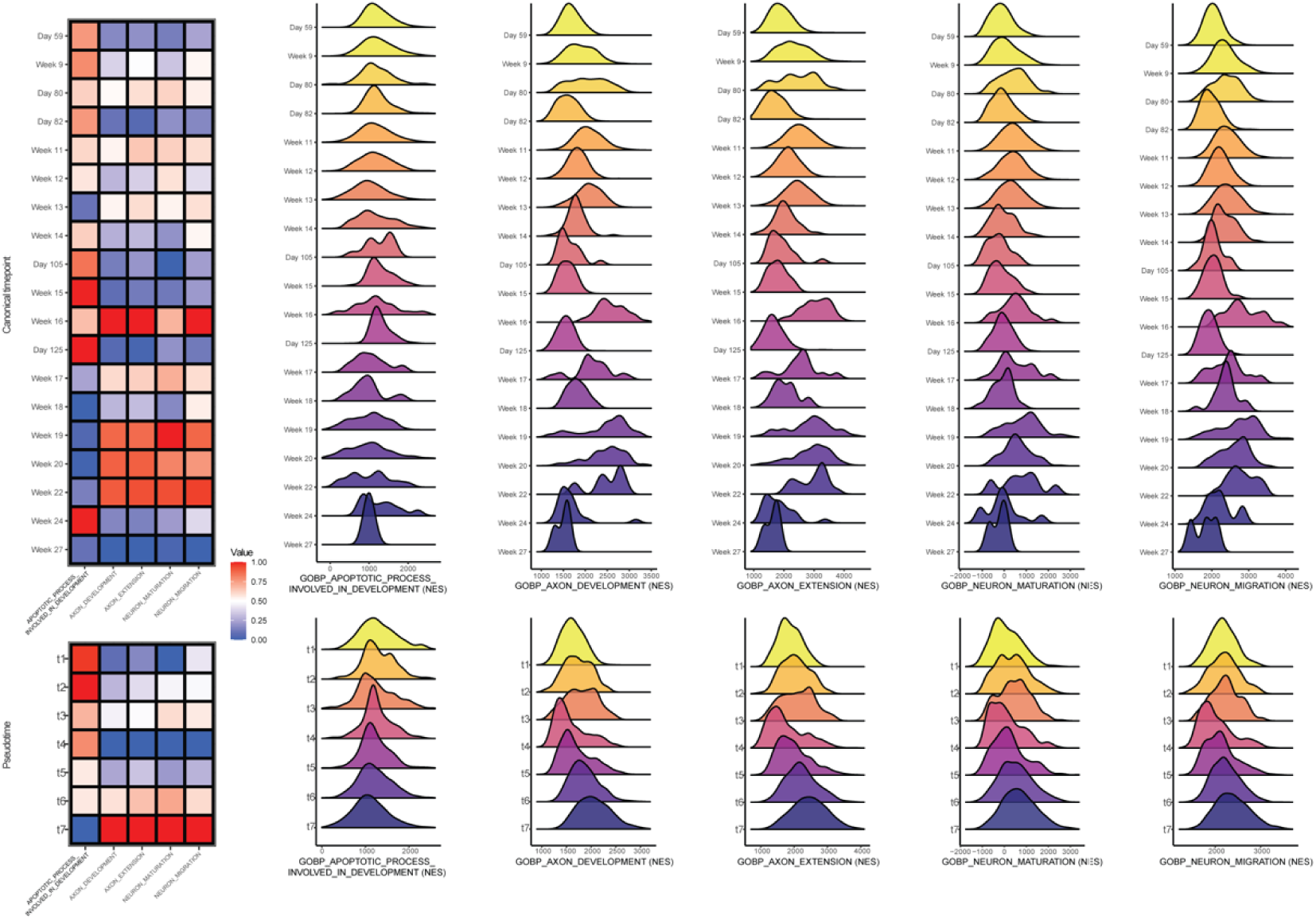
RGC pathway analysis during development. Cumulative double-normalized GSEA pathway analysis enrichment scores representing the timeframes for the pathway activity between canonical timepoint- and pseudotime-based approaches. Separated ridge enrichment plots comparing the pathway activity between canonical timepoint- and pseudotime-based approaches for GOBP_APOPTOTIC_PROCESS_INVOLVED_IN_DEVELOPMENT, GOBP_AXON_ DEVELOPMENT, GOBP_AXON_EXTENSION, GOBP_NEURON_ MATURATION, GOBP_NEURON_MIGRATION.

**Figure S7.**
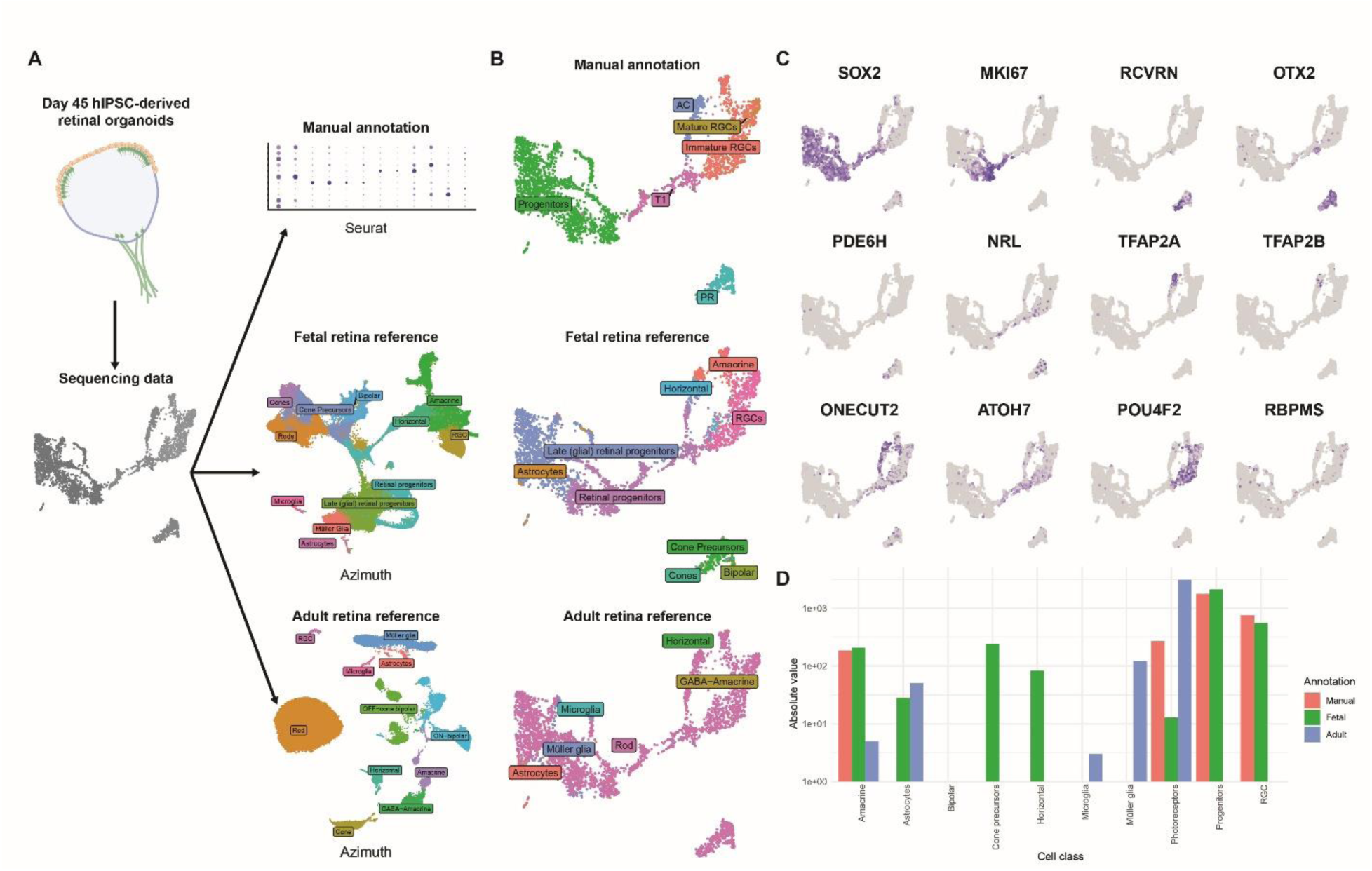
The comparison of annotation methods using sequencing data from human iPSC-derived retinal organoids at day 45 of differentiation. **(A)** Summarized pipeline for the annotation. The methods compared include manual annotation (with Seurat) and automated annotation (Azimuth-based) with human fetal retina atlas and adult retina reference. **(B)** Dimplots with annotation labels demonstrating the differences between the annotation approaches. **(C)** Exclusive markers for known retina cell classes presented in the feature plot format on the scRNAseq dataset of day 45 human iPSC-derived retinal organoids. **(D)** Barplot demonstrating the distribution of annotated cell classes and their frequencies upon the dataset annotation performed using three approaches: manual annotation, automated annotation using human fetal retina atlas reference, automated annotation using human adult retina reference.

**Table S1.** Software packages for single-cell RNA sequencing analysis.

| Software<br>/Package | Version | Link |
| --- | --- | --- |
| R | 4.2.1/4.3.1/4<br>.4.1 | https://cran.r-project.org/bin/ |
| RStudio | 2022.07.2-2023.06.1 | https://www.rstudio.com/products/rstudio/download/ |
| Azimuth | 0.4.6 | https://github.com/satijalab/azimuth |
| biomaRt | 2.52.0-2.56.1 | https://bioconductor.org/packages/release/bioc/html/biomaRt.html |
| CellChat | 1.5.0-2.1.0 | https://github.com/sqjin/CellChat |
| ComplexHeatmap | 2.16.0 | https://bioconductor.org/packages/release/bioc/html/ComplexHeatmap.html |
| cowplot | 1.1.1 | https://cran.r-project.org/web/packages/cowplot/ |
| CytoTRACE2 | 1.0.0 | https://github.com/digitalcytometry/cytotrace2 |
| dittoSeq | 1.8.1-1.12.2 | https://bioconductor.org/packages/release/bioc/html/dittoSeq.html |
| dplyr | 1.0.10-1.1.3 | https://dplyr.tidyverse.org/ |
| escape | 1.5.1-1.6.0 | https://github.com/ncborcherding/escape |
| future | 1.33.0 | https://cran.r-project.org/web/packages/future/index.html |
| ggalt | 0.4.0 | https://yonicd.github.io/ggalt/index.html |
| ggplot2 | 3.3.6-3.4.3 | https://ggplot2.tidyverse.org/ |
| GSEABase | 1.58.0-1.62.0 | https://bioconductor.org/packages/release/bioc/html/GSEABase.html |
| htmlwidgets | 1.5.4-1.6.2 | https://www.htmlwidgets.org/ |
| LRLoop | 0.1.0 | https://github.com/Pinlyu3/LRLoop |
| Matrix | 1.5-1-1.6-0 | https://cran.r-project.org/web/packages/Matrix/index.html |
| patchwork | 1.1.2-1.1.3 | https://patchwork.data-imaginist.com/ |
| plotly | 4.10.0 | https://plotly.com/r/ |
| reticulate | 1.26-1.32.0 | https://rstudio.github.io/reticulate/ |
| SCopeLoomR | 0.13.0 | https://github.com/aertslab/SCopeLoomR |
| sctransform | 0.4.0 | https://github.com/satijalab/sctransform |
| Seurat | 4.2.0-4.3.1 | https://satijalab.org/seurat/ |
| SeuratDisk | 0.0.0.9020 | https://github.com/mojaveazure/seurat-disk |
| Python | 3.7.4 | https://www.python.org/ |
| anndata | 0.8.0 | https://anndata.readthedocs.io/ |
| loompy | 3.0.7 | https://linnarssonlab.org/loompy/ |
| matplotlib | 3.5.3 | https://matplotlib.org/ |
| Mellon | 1.4.3 | https://github.com/settylab/Mellon |
| numpy | 1.21.6 | https://numpy.org/ |
| palantir | 1.2 | https://pypi.org/project/palantir/ |
| pandas | 1.3.5 | https://pandas.pydata.org/ |
| pySCENIC | 0.12.1 | https://pyscenic.readthedocs.io/ |
| scanpy | 1.9.3 | https://scanpy.readthedocs.io/ |
| scFates | 1.0.6 | https://scfates.readthedocs.io/en/latest/ |
| scvelo | 0.3.0 | https://scvelo.readthedocs.io/en/stable/index.html |
| seaborn | 0.12.2 | https://seaborn.pydata.org/ |

